# Thermally Tunable Heterodimeric Linkers Control Protein Tube Morphology

**DOI:** 10.1101/2025.11.24.689909

**Authors:** Masahiro Noji, Takuro Fujiwara, Yukihiko Sugita, Yuta Suzuki

## Abstract

Advances in artificial protein assembly design have enabled the construction of increasingly complex higher-order protein architectures. However, rationally tuning the morphology of such assemblies beyond their initially formed architectures remains challenging. Here, we show that the thermal responsiveness of a heterodimeric coiled-coil linker provides a molecular handle for tuning the formation and morphology of two-component protein tubes. Guided by computational stability predictions, we generated a panel of single-amino-acid variants in the M3L2/p66α linker and identified assembly-competent variants with distinct, experimentally accessible thermal transition behaviors. Incorporating two such variants into the tube-forming scaffold, together with the wild-type linker, enabled a focused comparison across three linker states, revealing stepwise shifts in tube-forming temperature windows and tube diameters. Notably, the variant with the lowest apparent thermal stability in this comparison uniquely accessed multi-walled tube architectures. Time-course electron microscopy of this variant revealed a sequential pathway from initially formed thin tubes to wall-thickened intermediates and ultimately to multi-walled states. Together, these findings establish linker thermal responsiveness as a design handle for tuning protein nanotube morphology and expanding the accessible structural states of designed protein tubes.

---

Higher-order protein assemblies support a wide range of biological functions by organizing protein subunits into defined yet adaptable architectures. Cytoskeletal filaments^1–4^, clathrin cages^5, 6^, viral capsids^7, 8^, and other natural assemblies^9, 10^ demonstrate that morphology is not simply an outcome of assembly, but a structural feature that can be tuned to regulate mechanical properties, spatial organization, and dynamic behavior. Recreating such morphological control in artificial protein assemblies would provide a route to protein nanostructures whose size, curvature, and architectural states can be modulated by molecular design^11–13^.

Recent breakthroughs in computational and modular protein design have enabled the de novo construction of increasingly sophisticated multi-component assemblies^14–18^. These approaches can specify interaction geometries with high precision, and newly reported quasisymmetric protein cage designs have elegantly demonstrated that curvature and related geometric parameters can be encoded into protein building blocks to program nanoscale size, symmetry, and morphology^19, 20^. These studies highlight the growing power of geometry-based design for creating target protein nanostructures with prescribed architectures. As a complementary direction, another opportunity is to tune the morphological states accessible from an established assembly framework. Protein nanotubes provide a useful platform for exploring this direction because their diameter, wall architecture, and assembly pathway can vary within the same overall tubular framework.

In our previous work, we developed two-component protein tubes assembled through the MBD3L2 (M3L2)/p66α heterodimeric coiled-coil linker^21^. These tubes exhibited a broad distribution of diameters and symmetries, indicating that the system can access multiple tubular morphologies from the same scaffold and linker framework. This observation suggested that the coiled-coil linker might provide a molecular handle for biasing the accessible tube morphologies. Because the M3L2/p66α linker serves as the central connecting element between the PuuE scaffold components, we hypothesized that altering its thermal responsiveness could shift the temperature window for tube formation, modulate tube diameter, and potentially change the structural states accessed during assembly.

To test this hypothesis, we generated a panel of single-point mutations in the M3L2/p66α coiled-coil linker, guided by computational stability predictions using ThermoMPNN^22^. Rather than aiming to redesign the overall scaffold geometry, this strategy was intended to perturb the thermal transition behavior of the connecting element while preserving heterodimer-mediated tube assembly. Biophysical characterization of the isolated coiled-coil variants identified assembly-competent linkers with distinct, experimentally accessible thermal transition behaviors. We then incorporated two representative variants into the PuuE tube-forming scaffold and compared them with the wild-type linker as three linker states. This focused comparison revealed stepwise shifts in the temperature window for tube formation and in tube diameter.

Notably, the variant with the lowest apparent thermal stability in this comparison uniquely accessed multi-walled tube architectures. Time-course electron microscopy further showed that these structures emerge through a sequential pathway from initially formed thin tubes to wall-thickened intermediates and ultimately to multi-walled states. Together, these results establish linker thermal responsiveness as a design handle for tuning protein nanotube morphology and expanding the structural states accessible from an established tube-forming framework.

## RESULTS AND DISCUSSION

### Engineering linker thermal responsiveness to tune protein tube morphology

To examine whether linker thermal responsiveness could tune protein tube morphology, we used our previously developed two-component tube system assembled through the MBD3L2 (M3L2)/p66α heterodimeric coiled-coil linker as a model platform (**Fig. 1**). In this system, PuuE scaffold components are connected through the M3L2/p66α linker, and the resulting tubes exhibit broad distributions of diameters and symmetries^21^. This morphological diversity suggested that the same scaffold and linker framework can access multiple tubular states, making it a suitable platform for testing whether linker properties can bias accessible tube morphologies. In our previous study, tube formation was promoted under conditions near the melting temperature (*T*_m_) of the M3L2/p66α interaction, suggesting that transient weakening and rearrangement of the linker can influence the tube morphologies accessed during assembly. Because p66α is intrinsically helical and structurally more defined, whereas M3L2 is more flexible and partially disordered^23, 24^, we focused our engineering efforts on M3L2 while leaving p66α unchanged to preserve the heterodimeric recognition mode. We reasoned that single-point mutations in M3L2 could shift the apparent thermal transition behavior of the coiled-coil linker and thereby bias the temperature window, diameter, and structural states accessed during tube assembly.

**Fig. 1.**
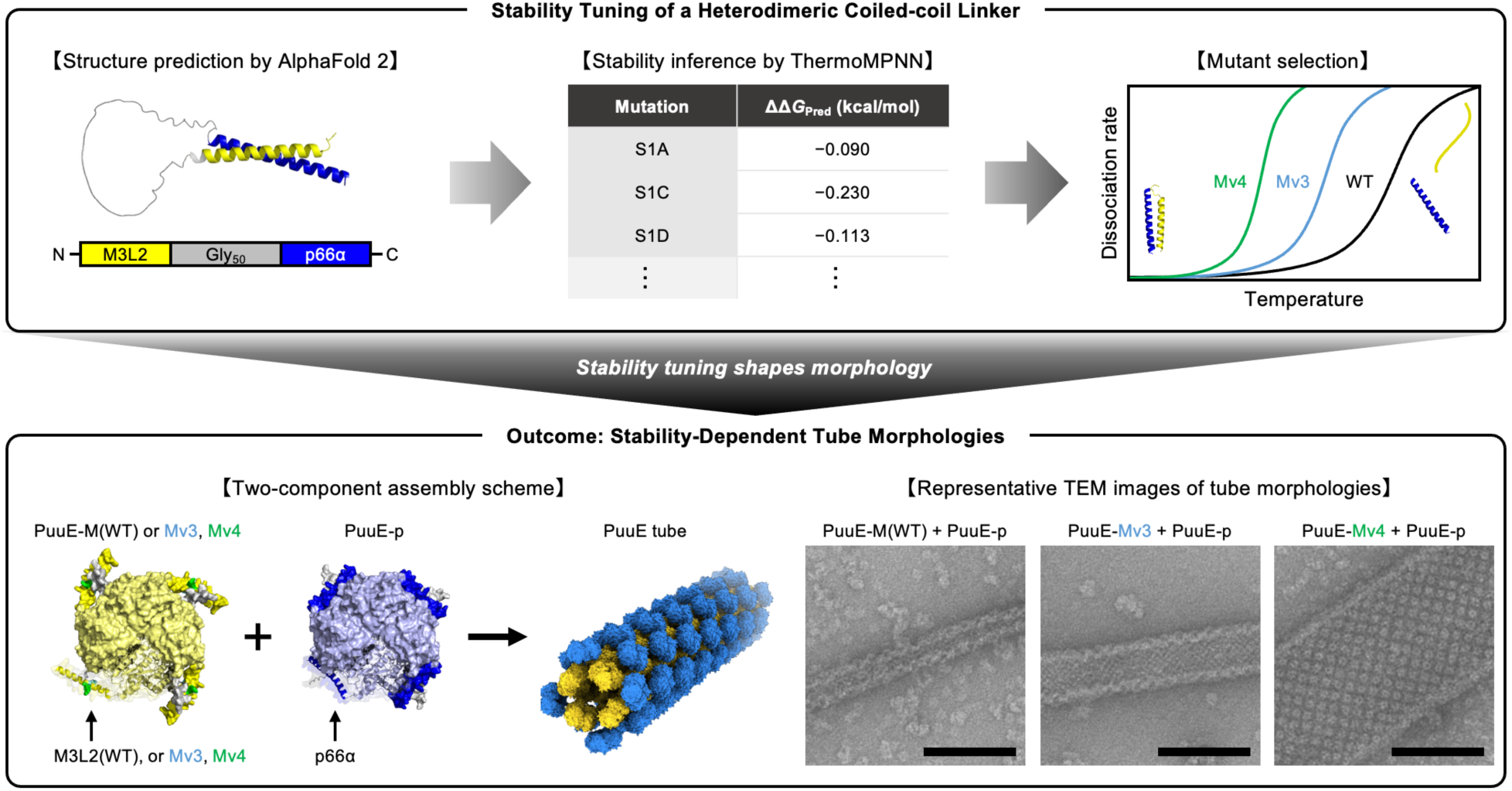
Thermal tuning of a heterodimeric coiled-coil linker modulates protein tube morphology. Building on our previously established PuuE-based two-component tube system, this study tests whether the thermal responsiveness of the M3L2/p66α linker can be used to bias accessible tube morphologies. Design workflow for generating stability-tunable M3L2/p66α coiled-coil variants using AF2 structural modelling, ΔΔ*G* predictions using ThermoMPNN, and selection of mutants spanning an engineered stability gradient (top). Assembly schematic and representative TEM images illustrating architecture-specific tube morphologies obtained with each variant (bottom). Stability-tuned M3L2 variants yield distinct tube morphologies —thin (WT), intermediate (Mv3), and thick (Mv4)—upon assembly with PuuE-p. Scale bar, 100 nm.

To generate M3L2 variants with altered thermal transition behavior, we used ThermoMPNN to guide single-point mutagenesis of the M3L2/p66α coiled-coil linker. ThermoMPNN predicts mutation-induced changes in protein stability (ΔΔ*G*_Pred_, defined here as the predicted change in coiled-coil stability upon mutation)^22^. Because ThermoMPNN is not directly applicable to multimeric complexes, we first generated an M3L2-Gly_50_-p66α heterodimer model using AlphaFold 2 (AF2)^25^ and used the highest-confidence model as the input structure for stability inference (**Fig. 1**). We then restricted mutation scanning to previously reported key residues within the M3L2^24^ and selected 13 single-point mutations with destabilizing ΔΔ*G*_Pred_ values (**Fig. 2a**, **Table 1, and Supplementary Table 1**). This strategy used computational stability prediction in an inverse manner: predicted destabilization was used not to optimize linker stability or directly predict tube morphology, but to identify experimentally tractable perturbations of linker thermal behavior. This direction was motivated by our previous finding that tube formation was promoted near the M3L2/p66α melting temperature, suggesting that partial weakening of the linker can facilitate productive rearrangements during assembly. We therefore reasoned that modest linker destabilization could shift this dynamic assembly regime toward lower, protein-compatible temperatures while preserving heterodimer-mediated tube formation.

**Fig. 2.**
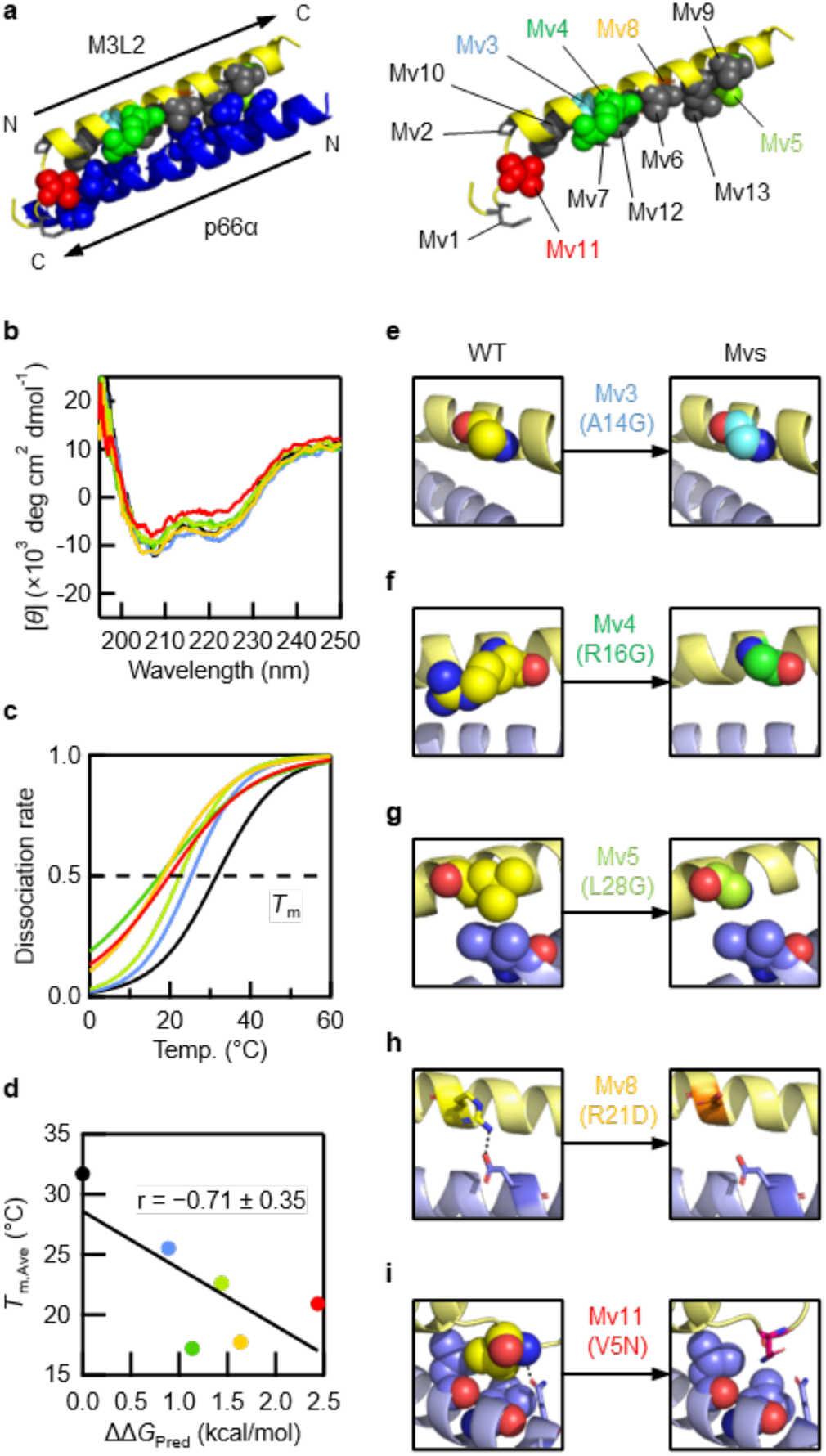
Stability-guided linker engineering identifies variants with distinct thermal transition behaviors. **a,** Schematic representations of the M3L2/p66α heterodimeric coiled-coil model generated by AF3 (left) and the targeted mutation sites (right). **b,** CD spectra of WT and selected M3L2 mutants complexed with p66α indicating preserved coiled-coil secondary structure. **c,** Thermal denaturation profiles of WT and selected M3L2 mutants. **d,** Correlation between predicted ΔΔ*G*_Pred_ values using ThermoMPNN and the experimentally determined *T*_m,Ave_. **e**–**i,** AF3 structural models illustrating mutation-dependent local perturbations within the M3L2/p66α interface. Mutated residues are shown in space-filling (ball) representation to highlight steric volume and hydrophobic packing effects, whereas neighbouring residues involved in hydrogen-bonding or other directional interactions are shown in stick representation highlighting how each substitution alters the local interaction environment.

**Table 1.** M3L2 wild-type and selected mutants. Predicted stability changes for point mutations (ΔΔ*G*_Pred_) obtained using ThermoMPNN are shown for M3L2 wild-type (WT) and its selected mutants (Mvs). Average melting temperatures (*T*_m,Ave_) determined from CD measurements are provided for coiled-coil complexes formed between p66α and each variant. Source data are provided as a Source Data file.

| Name | Mutation | $\Delta\Delta G_{\text{Pred}}$ (kcal/mol) | $T_{\text{m,Ave}}$ (°C) |
| --- | --- | --- | --- |
| WT | None | 0 | 31.7 |
| Mv3 | A14G | 0.888 | 25.5 |
| Mv4 | R16G | 1.136 | 17.2 |
| Mv5 | L28G | 1.443 | 22.6 |
| Mv8 | R21D | 1.637 | 17.7 |
| Mv11 | V5N | 2.439 | 20.9 |

We next examined how the selected M3L2 mutations affected heterodimer formation with p66α and the thermal behavior of the resulting coiled-coil complexes. To characterize these effects independently of the PuuE scaffold, we constructed N-terminal His-tagged versions of M3L2 (WT and all mutants) and p66α for biophysical analysis in isolation and in complex. Both M3L2 and p66α were expressed in soluble form in *E. coli*, but 6×His-TEVcs-M3L2(WT) was recovered inefficiently by nickel-affinity chromatography because much of the protein appeared in the flowthrough. Extending the tag to 8×His-TEVcs enabled robust purification of M3L2(WT), and all 13 M3L2 variants were therefore constructed and purified in the same 8×His-tagged format to ensure consistency across the peptide dataset. To assess coiled-coil formation between M3L2 and p66α, we performed circular dichroism (CD) spectroscopy using the purified His-tagged peptides (**Fig. 2b and Supplementary Fig. 1**). Individually, M3L2(WT) exhibited a largely disordered conformation, whereas p66α adopted a more helical structure as expected. Mixing the two peptides increased the α-helical signal and produced a temperature-dependent unfolding transition, consistent with reconstitution of the M3L2/p66α heterodimeric coiled-coil. The averaged melting temperature (*T*_m, Ave_) of the reconstituted WT complex was 31.7 °C (**Fig. 2c, Supplementary Fig. 2, Table 1, and Supplementary Table 1**), close to the previously reported value of 34.8 °C for the M3L2/p66α coiled-coil^24^, despite minor sequence differences outside the core coiled-coil region. These results confirmed that the purified peptide system preserves the essential thermal behavior of the heterodimer and is suitable for evaluating the designed M3L2 variants.

Applying the same CD and thermal unfolding analyses to the 13 engineered M3L2 variants showed that several mutations reduced or disrupted well-defined coiled-coil formation with p66α, consistent with substantial perturbation of the heterodimeric interaction (**Fig. 2b and Supplementary Fig. 1**). Among the full mutant panel, five variants—Mv3, Mv4, Mv5, Mv8, and Mv11—formed M3L2/p66α complexes with sufficiently well-defined CD profiles to permit reliable thermal denaturation analysis. Thermal unfolding experiments on the five variants with well-defined CD profiles yielded measurable *T*_m_ values that were lower than that of the WT complex and spanned a range of apparent thermal stabilities (**Fig. 2c, Supplementary Fig. 2, Table 1, and Supplementary Table 1**). Across this experimentally tractable subset, the averaged melting temperatures (*T*_m,Ave_) showed a negative correlation with the ThermoMPNN-predicted ΔΔ*G*_Pred_ values (r = −0.71 ± 0.35), consistent with the intended direction of linker destabilization (**Fig. 2d**). The remaining variants showed weak CD signatures and poorly defined thermal denaturation profiles, preventing reliable *T*_m_ determination (**Supplementary Figs. 1 and 2**). Thus, the mutation panel identified a subset of M3L2 variants with distinct apparent thermal stabilities while also defining variants in which the M3L2/p66α interaction was too strongly perturbed for reliable thermal analysis. These results provided a biophysical basis for identifying representative linker variants that retained measurable coiled-coil formation while spanning the experimentally accessible range of linker thermal behavior for subsequent assembly studies. This distinction was important for selecting variants that perturbed the M3L2/p66α interaction while preserving sufficient heterodimeric character for subsequent assembly studies.

### Structural evaluation of engineered coiled-coil variants

To further evaluate how the selected mutations might affect the M3L2/p66α interface, we generated AlphaFold 3 (AF3)^26^ models for each M3L2/p66α heterodimer pair (**Fig. 2e–i and Supplementary Fig. 3**). These models were used as qualitative structural guides to assess whether each substitution was likely to preserve, weaken, or disrupt the local coiled-coil interaction environment. Thus, the AF3 analysis was used to complement the CD and thermal unfolding data rather than to provide quantitative stability estimates.

The AF3 models provided structural context for the different experimental behaviors observed across the mutant panel. For Mv1 (Q3Y), which had only a small, predicted destabilizing ΔΔ*G*_Pred_ value, the model suggested disruption of an interchain hydrogen bond near the flexible N-terminal region of M3L2 (**Supplementary Fig. 3a**). This local environment may not be fully captured by stability prediction alone, particularly because the N-terminal region contains partially disordered character. Mv2 (P7G) also showed weaker coiled-coil behavior than expected from its predicted ΔΔ*G*_Pred_ value, suggesting that replacement of the helix N-cap proline with glycine can influence local helix stability^27, 28^ in ways not fully captured by the stability prediction (**Supplementary Fig. 3b**).

In contrast, the AF3 models of Mv3 and Mv4 suggested that these mutations do not directly abolish the overall M3L2/p66α recognition interface (**Fig. 2e, f**). This feature made them suitable candidates for subsequent incorporation into the PuuE tube-forming scaffold, because they were expected to perturb linker thermal behavior while retaining the heterodimeric interaction required for assembly. Importantly, Mv3 and Mv4 also displayed distinct, experimentally measurable thermal transition behaviors, providing a practical pair for testing how different linker states influence tube formation and morphology. Notably, although both variants introduce glycine substitutions, the mutations occur in different local structural environments. This difference provides a structural basis for anticipating that Mv3 and Mv4 could perturb linker dynamics to different extents after incorporation into the tube-forming scaffold.

For variants with larger predicted destabilizing effects, including Mv5–Mv13, AF3 models generally suggested perturbation of interchain packing interactions (**Fig. 2g–i, Supplementary Fig. 3c–h, Table 1, and Supplementary Table 1**). These models were useful for identifying substitutions likely to alter the local coiled-coil interaction environment, although the extent of structural perturbation inferred from the models did not always map directly onto the CD signal or the ability to obtain reliable thermal unfolding profiles. Taken together, the CD spectra, thermal unfolding profiles, and AF3 models provided a basis for selecting variants that retained sufficient heterodimeric character while spanning distinct apparent thermal stabilities. On this basis, Mv3 and Mv4 were selected as representative assembly-competent linkers for incorporation into the two-component PuuE tube system. This integrated selection strategy helped separate variants that primarily disrupted measurable coiled-coil behavior from variants that preserved the interaction while altering linker thermal responsiveness, which was essential for testing morphology tuning in the assembled tube system.

### Effect of linker thermal responsiveness on two-component tube formation

Having identified Mv3 and Mv4 as assembly-competent variants with distinct apparent thermal stabilities, we next introduced these linker mutations into the PuuE tube-forming scaffold to test how linker thermal responsiveness affects two-component tube formation (**Fig. 3a**). Together with the wild-type M3L2/p66α linker, Mv3 and Mv4 provided three experimentally tractable linker states for comparing tube formation, temperature dependence, tube diameter, and the structural states accessed during assembly. This design allowed us to evaluate whether perturbing the thermal behavior of the connecting element, rather than redesigning the scaffold architecture or introducing new assembly interfaces, could bias the morphology of the resulting protein tubes.

**Fig. 3.**
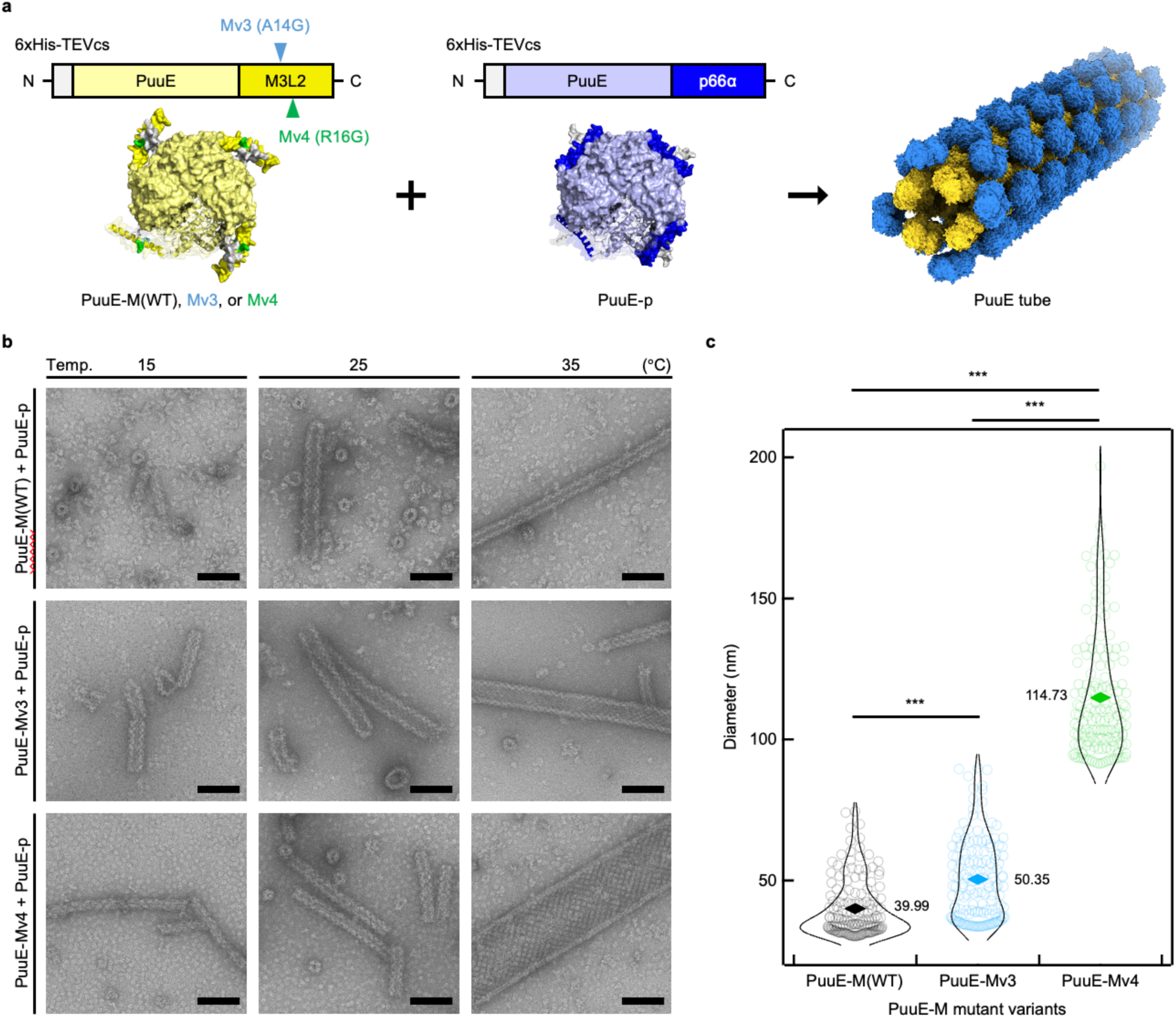
Linker thermal responsiveness shifts the temperature dependence and morphology of two-component protein tubes. **a,** Schematic representation of the two-component tube-forming system, showing PuuE-M(WT) or its stability-tuned variants (PuuE-Mv3, PuuE-Mv4) paired with PuuE-p via the C-terminal M3L2/p66α heterodimeric linker. **b,** Representative nsTEM images of tubes formed by PuuE-M(WT), PuuE-Mv3, and PuuE-Mv4 in complex with PuuE-p at 15, 25, and 35 °C. Additional representative TEM images across temperatures are provided in **Supplementary** Fig. 5. **c,** Quantified tube diameter distributions for PuuE-M(WT), PuuE-Mv3, and PuuE-Mv4 after 24 h of incubation at 35 °C. Mean values are shown with diamond-shaped markers. \*\*\**p* < 0.001 (*p*-values; WT–Mv3 = 4.21 × 10^−12^, Mv3–Mv4 = 8.87 × 10^−88^, and WT–Mv4 = 2.11 × 10^−7^, two-sided Welch’s *t* test). Scale bar, 100 nm.

As reported previously^21^, PuuE-M(WT) and PuuE-p form two-component tubes through the C-terminal M3L2/p66α coiled-coil linker, with efficient tube formation near 35 °C (**Fig. 3b**). We introduced the Mv3 and Mv4 mutations individually into PuuE-M(WT) to generate PuuE-Mv3 and PuuE-Mv4 and first examined whether these variants retained the ability to assemble with PuuE-p (**Fig. 3a**). Both PuuE-Mv3 and PuuE-Mv4 were purified successfully and did not form higher-order structures when examined alone by negative-stain transmission electron microscopy (nsTEM; **Supplementary Fig. 4**). Upon mixing with PuuE-p at 35 °C, both variants formed tubular assemblies, confirming that the Mv3 and Mv4 linkers preserve the heterodimer-mediated assembly competence required for two-component tube formation (**Fig. 3b and Supplementary Fig. 5**). Having established tube formation across PuuE-M(WT), PuuE-Mv3, and PuuE-Mv4, we next compared how the three linker states affect the temperature window for tube formation and the morphology of the resulting tubes.

To evaluate how linker thermal responsiveness affects the temperature window for tube formation, we compared PuuE-M(WT), PuuE-Mv3, and PuuE-Mv4 across different incubation temperatures. As reported previously^21^, the PuuE-M(WT)/PuuE-p system exhibits temperature dependence, with efficient tube formation near 35 °C and limited assembly at lower temperatures. Consistent with this behavior, mixtures of PuuE-M(WT) and PuuE-p formed micron-scale tubes efficiently at 35 °C, whereas only sparse tube formation was observed at 15 °C (**Fig. 3b and Supplementary Fig. 5**). In contrast, the Mv3 and Mv4 variants shifted tube-forming behavior toward lower temperatures. At 15 °C, PuuE-Mv3 showed partial tube formation, whereas PuuE-Mv4 retained robust tube-forming capability and produced discernible micron-scale tubes under conditions where the WT system showed little assembly. Thus, across the three linker states, decreasing apparent thermal stability was associated with a stepwise expansion of the tube-forming temperature window toward lower temperatures. This trend suggests that weakening the linker does not simply disrupt assembly but instead shifts the range of conditions under which productive tube formation can occur.

Having observed that the WT, Mv3, and Mv4 linkers differ in their tube-forming temperature windows, we next asked whether these linker variants also alter the physical morphology of the resulting tubes. Because our previous work showed that PuuE-based tubes can access a broad range of diameters^21^, we quantified tube diameters for PuuE-M(WT), PuuE-Mv3, and PuuE-Mv4 assembled with PuuE-p at 35 °C. Quantitative diameter analysis showed a stepwise shift across the three constructs, with PuuE-M(WT) producing the smallest-diameter tubes, PuuE-Mv3 yielding intermediate diameters, and PuuE-Mv4 forming the largest-diameter tubes (**Fig. 3b, c and Supplementary Fig. 5**). These results show that single-amino-acid changes in the connecting M3L2/p66α linker can bias the diameter distribution of two-component protein tubes by altering linker thermal responsiveness. Together, the temperature-window and diameter analyses indicate that local perturbation of the linker is sufficient to redirect both when tubes form and which tubular morphologies are preferentially accessed within the same assembly framework, providing a route toward more precise access to distinct tubular arrangements in artificial protein assemblies.

### Formation process and structural features of multi-walled tube architectures

In addition to the diameter increase described above, PuuE-Mv4 tubes formed at 35 °C displayed multi-walled tube architectures that were not apparent in the WT or Mv3 samples **(Fig. 4a, b and Supplementary Fig. 6a**). Cryo-electron tomography (cryo-ET) further supported this assignment, with some assemblies containing up to four tube layers across multiple fields of view (**Supplementary Fig. 7a–d**). These observations indicate that multi-walled tube formation is a reproducible morphological outcome of the Mv4-containing system under these assembly conditions. The multi-walled structures were often distorted or incomplete in regions of thin ice, including flattened tubes and assemblies in which the outer layer only partially covered the inner tube or was locally discontinuous (**Supplementary Fig. 7e**). These features suggest that the multi-walled tubes are flexible and susceptible to deformation during cryo-EM sample preparation. Although the observed structures were not always idealized multi-layered tubes, the combined nsTEM and cryo-ET observations support the conclusion that Mv4 uniquely accesses multi-walled tube states under these conditions.

**Fig. 4.**
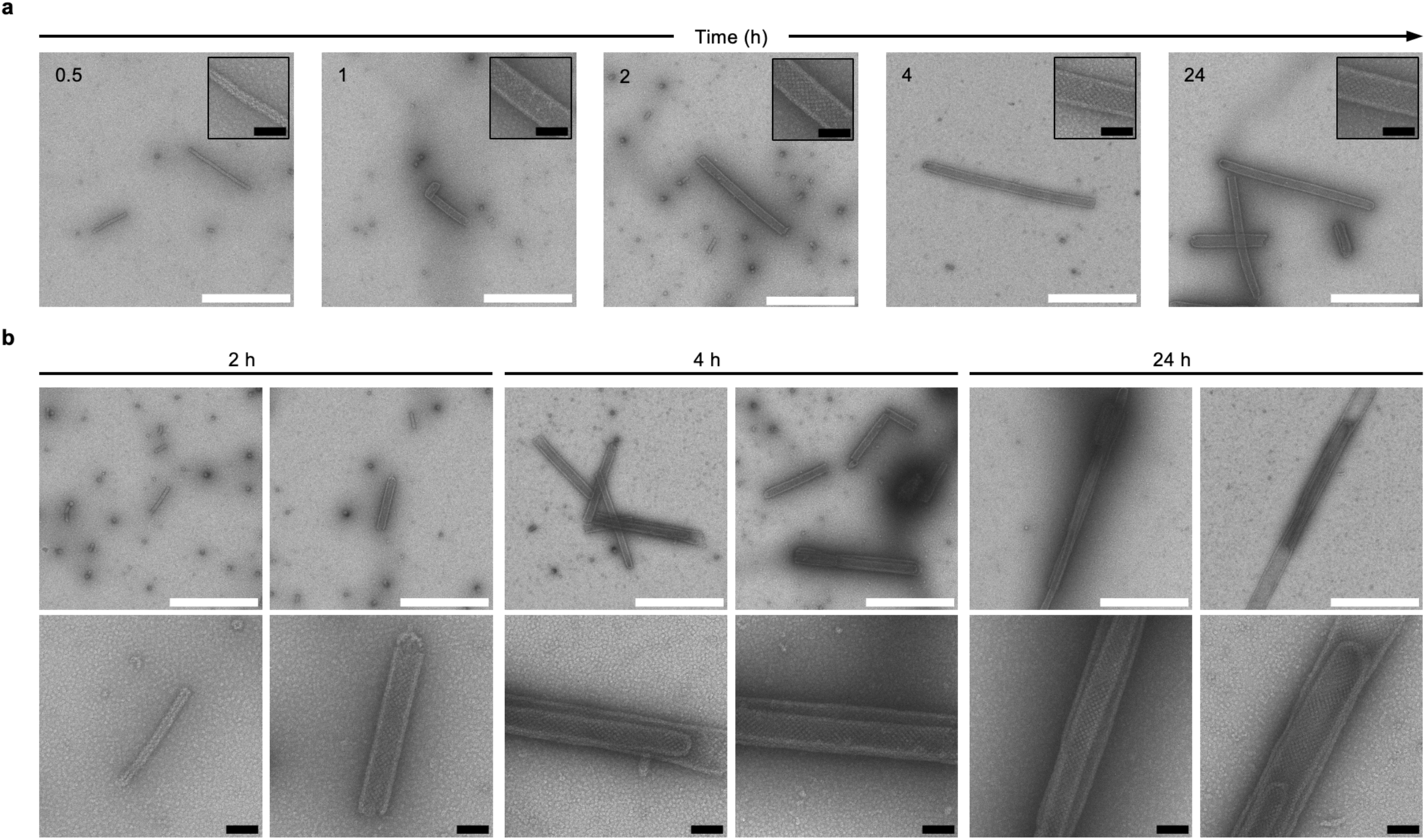
PuuE-Mv4 accesses multi-walled tube states through a stepwise assembly process. **a,** Time-resolved nsTEM images of PuuE-Mv4 assembled with PuuE-p at 35 °C, showing progressive elongation of tubes over time (0.5, 1, 2, 4, and 24 h). Insets show magnified views of representative tube segments at each time point. **b,** Representative nsTEM images comparing tubes lacking tube-in-tube features (2 h) with those in which tube-in-tube architectures begin to appear (4 h) and fully develop (24 h). Additional time points and further representative images are provided in **Supplementary** Fig. 6. Scale bars: white, 1 µm; black, 100 nm.

To examine how these Mv4-specific multi-walled tube states arise during assembly, we performed time-resolved nsTEM imaging of PuuE-Mv4/PuuE-p mixtures incubated at 35 °C and analyzed the evolution of tube morphology over time (**Fig. 4 and Supplementary Fig. 6a**). At early time points (0–30 min), PuuE-Mv4/PuuE-p mixtures primarily contained thin, single-walled tubes, typically several hundred nanometers in length. After 1–2 h, thicker tubes became more prominent, indicating that tube growth and wall thickening continued after initial tube formation. Multi-walled tube architectures were not apparent at these early stages. By approximately 4 h, however, multi-walled structures began to appear, including tubes in which an additional tubular layer was partially or fully present within a pre-existing tube. These multi-walled structures persisted from 4 h to 24 h, suggesting that they represent a later-formed structural state rather than an early assembly intermediate. This temporal order argues against simultaneous multilayer formation and instead supports a stepwise pathway in which wall thickening precedes the appearance of multi-walled tube states.

In addition to these multi-walled states, nsTEM revealed local structural irregularities in PuuE-Mv4, including incomplete or misaligned tube junctions and displaced wall segments (**Supplementary Fig. 6b**). These features were not apparent in the WT or Mv3 samples under the same conditions, suggesting that they are associated with the Mv4-specific pathway toward multi-walled tube formation. Although the images do not directly establish a causal role, these local irregularities are consistent with transient growth states that could contribute to wall thickening or secondary layer formation during assembly. In this view, the reduced apparent thermal stability of the Mv4 linker may increase the probability of local rearrangements during growth, allowing imperfect tube configurations to be reorganized into thicker or multi-walled structures. Together, the time-course observations support a sequential assembly pathway in which PuuE-Mv4 first forms thin single-walled tubes, then wall-thickened tubes, and finally multi-walled tube states. Thus, linker thermal responsiveness not only shifts the temperature window and diameter of tube formation but also changes the structural states accessible during assembly. These findings extend the role of linker thermal responsiveness from modulating tube formation and diameter to controlling access to alternative tube architectures within the same two-component assembly framework.

## Conclusion

This work shows that linker thermal responsiveness can serve as a molecular handle for tuning the morphology of two-component protein tubes. Guided by computational stability predictions and biophysical characterization, we identified M3L2 variants that retained heterodimeric assembly competence while displaying distinct apparent thermal stabilities. Incorporation of two representative variants into the PuuE tube-forming scaffold, together with the wild-type linker, revealed that single-amino-acid changes in the connecting linker can shift the tube-forming temperature window, bias tube diameter distributions, and enable access to multi-walled tube states. These effects were achieved by modulating the M3L2/p66α linker without requiring redesign of the overall scaffold architecture or introduction of new assembly interfaces. This strategy complements recent geometry-based protein nanostructure design approaches, in which curvature, symmetry, or building-block geometry can be programmed to generate prescribed target architectures^14, 15, 17–20^. The present work adds to this design landscape by showing that local modulation of linker behavior can tune the structural states accessible from an established tube-forming framework. More broadly, it suggests that dynamic interaction modules can serve as design elements for expanding the morphological space of engineered protein assemblies.

## Methods

### Computational engineering of the M3L2/p66α coiled-coil interaction

The structure of M3L2-Gly_50_-p66α was predicted using AlphaFold 2.3 (DeepMind)^25^. The most reliable model (ranked_0 model) was used as the input for ThermoMPNN^22^ to infer ΔΔ*G*_Pred_ values for all possible single-mutations. The resulting ΔΔ*G*_Pred_ scores were examined, and for the previously reported coiled-coil key residues in M3L2, mutations with the highest positive ΔΔ*G*_Pred_ values were selected. The overall computational workflow is summarized in **Fig. 1**, and the selected mutations are listed in **Table 1** and **Supplementary Table 1**. Amino acid sequences for all constructs used in this study are provided in **Supplementary Table 2**.

### Plasmids and cloning

The plasmids encoding N-terminal 6×His-TEVcs-tagged PuuE-M(WT) and PuuE-p were previously constructed. The plasmids encoding N-terminal 6×His-TEVcs-tagged M3L2(WT) and p66α were generated from the PuuE-M(WT) and PuuE-p plasmids, respectively, by removal of the PuuE coding region. The plasmid encoding N-terminal 8×His-TEVcs-tagged M3L2(WT) was generated from the 6×His-TEVcs-M3L2(WT) plasmid by introducing two additional His codons. The plasmids encoding N-terminal 8×His-TEVcs-tagged Mv series (Mvs) and 6×His-TEVcs-tagged PuuE-Mv series (PuuE-Mvs) were generated from the 8×His-TEVcs-M3L2(WT) and PuuE-M(WT) plasmids, respectively, by site-directed substitution of the corresponding codons. All primers used in this study were purchased from Eurofins Genomics, and all primer sequences are shown in **Supplementary Data 1**. Mutagenesis was performed using the KOD - Plus-Mutagenesis Kit (TOYOBO) according to the manufacturer’s instructions. The 6×His-TEVcs-M3L2(WT), 8×His-TEVcs-M3L2(WT), Mvs, and p66α plasmids were amplified in *E. coli* strain BL21 (DE3) derived ECOS SONIC competent cells (NIPPON GENE) and extracted using the NucleoSpin Plasmid EasyPure (MACHEREY-NAGEL) according to the manufacturer’s protocol. The PuuE-Mvs plasmids were amplified in *E. coli* strain DH5α (NIPPON GENE) and purified using the same procedure. All construct sequences were verified by Eurofins Genomics. Amino acid sequences for all constructs used in this study are provided in **Supplementary Table 2**.

### Protein expression and purification

The recombinant proteins of N-terminal 6× or 8×His-TEVcs-tagged M3L2(WT), Mvs, and p66α were expressed using *E. coli* strain BL21 (DE3) derived ECOS SONIC competent cells as follows. Glycerol stocks of transformed cells were used to inoculate 5 mL of liquid LB broth supplemented with 100 μg/mL ampicillin (Amp) and grown overnight at 37 °C and 200 rpm. Overnight cultures were diluted into 1 L of LB-Amp broth and grown at 37 °C and 200 rpm until reaching an optical density at 600 nm of 0.4–0.6. Protein expression was induced by adding 0.1 mM isopropyl-β-D-thiogalactopyranoside, and the cultures were grown at 16 °C for 18–20 h. Cells were harvested by centrifugation at 15,317 × *g* for 5 min at 4 °C and stored at –80 °C.

The recombinant proteins of PuuE-M(WT), PuuE-Mvs, and PuuE-p were expressed using *E. coli* strain BL21 (DE3) (NIPPON GENE) co-transformed with a pGro7 chaperone plasmid (Takara Bio). After transformation, colonies grown overnight on LB agar plates supplemented with Amp and 20 μg/mL chloramphenicol (Crm) at 37 °C were used to inoculate 5 mL of liquid LB-Amp-Crm broth and cultured overnight at 37 °C and 200 rpm. Overnight cultures were diluted into 1 L of liquid LB-Amp-Crm broth supplemented with 0.5 mg/mL L-arabinose and grown at 37 °C and 200 rpm until reaching an optical density at 600 nm of 0.4–0.6. Protein induction and cells harvest was performed as described above.

All recombinant proteins were purified as follows. Cell pellets were thawed at 37 °C, resuspended in 60 mL of ice-cold purification buffer (20 mM Tris-HCl, pH 8.0, containing 300 mM NaCl), and lysed by sonication (9 min, 1:2 on/off cycles, 70% amplitude; SFX250, Branson) on ice. Cell debris was removed by centrifugation at 15,317 × *g* for 30 min at 4 °C. The supernatants were passed through a 0.45-μm pore size membrane filter (Merck), loaded onto a HisTrap FF crude column (Cytiva) pre-equilibrated with the purification buffer, and washed with 5 column volumes of 2% elution buffer (20 mM Tris-HCl, pH 8.0, containing 300 mM NaCl and 1 M imidazole; 2% means 20 mM imidazole). Target proteins were eluted with 10 column volumes of elution buffer with a linear gradient of 2–40% (i.e., 20–400 mM imidazole). Fractions containing the target proteins as verified by UV absorbance and SDS-PAGE were dialysed twice against a 50-fold volume of experimental buffer (50 mM Tris-HCl, pH 8.0, containing 100 mM NaCl and 0.5 mM EDTA) at 4 °C. Purified proteins were concentrated by Amicon Ultra centrifugal filter unit (Merck) with an appropriate molecular weight cutoff, followed by filtration through a 0.45 μm pore size membrane filter (Merck). Protein concentrations were determined by measuring absorbance at 280 nm using a NanoDrop OneC spectrophotometer (Thermo Scientific). The molar extinction coefficients at 280 nm for the proteins were calculated based on amino acid composition^29^. The concentrated proteins were flash-frozen in liquid nitrogen and stored at –80 °C until use.

### Circular dichroism (CD) spectrum measurements

All proteins were thawed immediately before CD measurements and kept on ice. Each sample was prepared in a 1.5-mL microtube using experimental buffer to adjust the final protein concentration to 5.0 µM for single-component samples or 2.5 + 2.5 μM for binary complexes, in a total volume of 200 μL at 25 ± 1 °C. Far-UV CD spectra were recorded at a wavelength of 195 to 250 nm using a J-1100 spectropolarimeter (JASCO) equipped with a 1-mm-path-length quartz cuvette, at the temperatures indicated in the manuscript.

Thermal denaturation measurements were performed over a temperatures range of 0 to 60 °C, with a heating rate of 1 °C/min, while monitoring ellipticity at 222 nm. All spectra data were collected using Spectra Manager version 2.5 (JASCO) and expressed as mean residue ellipticity. The *T*_m_ value of each sample was determined from the thermal denaturation curve by sigmoid fitting using Igor Pro 9 version 9.0.5.1 (WaveMetrics).

### Tube formation

All proteins were thawed immediately before tube formation experiments and kept on ice. Each sample was prepared in a 1.5-mL microtube using experimental buffer to adjust the concentration to 12.5 μM of each protein (considering tetramer equivalence), with a final volume of 100 μL at 25 ± 1 °C. Sample incubation was performed using a ThermoMixer C (Eppendorf) or a MATRIX Orbital Delta Plus (IKA) with shaking at 300 rpm, at the temperatures described in the manuscript.

### Negative-stain transmission electron microscopy (nsTEM)

A naked G600TT copper grid (Nisshin EM) was carbon-coated using a VE-2030 (VACUUM DEVICE). The grid was glow-discharged using a PIB-10 (VACUUM DEVICE). Then, a 5-μL aliquot of the sample solution was placed on the grid for 1 min, and the remaining solution was removed with filter paper (No. 2, ADVANTEC) followed by rinsing thrice with a 5-μL aliquot of Milli-Q water. After blotting off the water with filter paper, the sample was stained briefly with a 3-μL aliquot of 2% (w/v) uranyl acetate solution three times. The remaining solution was removed with filter paper and the grid was dried on the benchtop. TEM observation was performed using a transmission electron microscope HT-7700 (Hitachi) with an acceleration voltage of 80 kV. The images were recorded using HT-7700 control software version 02.22 (Hitachi).

### Cryo-electron tomography (Cryo-ET)

The sample stock solution containing 12.5 µM of each protein was diluted 10-fold, and a 2.7-µL aliquot of the diluted specimen solution was applied onto glow-discharged Quantifoil R1.2/1.3 Gold 200 mesh grids (Quantifoil Micro Tools GmbH). After blotting excess solution from the grids with filter paper, specimens were rapidly frozen in liquid ethane on a Vitrobot Mark IV (Thermo Fisher Scientific). Cryo-ET datasets were collected using a Glacios cryo-TEM (Thermo Fisher Scientific) operated at an acceleration voltage of 200 kV, equipped with a Falcon4 direct electron detector at the Institute for Life and Medical Sciences, Kyoto University. Tilt-series were collected with a dose-symmetric tilt scheme^30^ using TEM Tomography 5 software (Thermo Fisher Scientific). The tilt span of ±60° was used in 3° increments, starting at 0^°^. All tilt-series were collected in nanoprobe mode with a defocus of -4.0 µm and TIFF compression enabled. Data were acquired at a dose rate of 10.6 e^−^/pixel/s, a pixel size of 1.55 Å, and a total exposure of 144 e^−^/Å^2^ per tilt series. Eucentric height estimation was performed once for each tilt angle using stage-tilt method in TEM Tomography 5 software. Regions of interest were added manually, and positions were saved. Tracking and focusing were performed before and after each tilt step. All data were processed using RELION-5 tomography pipeline^31^. TIFF movie frames were motion corrected using RELION’s own implementation and gain normalization using a gain reference image generated by the “relion_estimate_gain” command in RELION-5. Tilt-series alignment was performed using AreTomo2^32^, which simultaneously estimates the contrast transfer function (CTF) for each tilt image during alignment. The aligned tilt series were subsequently used to reconstruct tomograms. All tomographic reconstructions were binned to yield a final pixel size of 12.4 Å, and were visualized using IMOD software package^33^.

### Tube diameter analysis

For each sample incubated at 35 °C, more than 200 clearly discernible tubes were picked up manually on 10k-magnification nsTEM images. Tube segments in which tubes overlapped were excluded from analysis. Tube diameters were measured using ImageJ (Fiji) version 1.53^34^. For quantitative comparison, a representative subset of 150 tubes was selected from the measurable population using the same inclusion criteria across all samples, focusing on the larger-diameter species that predominated under the tube-forming conditions. Diameter distributions and plots were generated using Igor Pro 9.

### Molecular modelling

The structure of M3L2-Gly_50_-p66α, PuuE-M(WT), and PuuE-p was predicted by AlphaFold 2.3 (DeepMind)^25, 35^. All other predicted protein structures were generated using AlphaFold 3 (DeepMind)^26^. Cartoon representations of protein models were drawn using PyMOL version 2.5 (Schrödinger)^36^ and UCSF ChimeraX version 1.9 (UCSF RBVI and NIH)^37^.

### Statistics and Reproducibility

For the analysis of tube diameter, a two-sided Welch’s *t* test was performed to test the null hypothesis, and the exact *p* values are provided in the figure legend. Each of the tube formation experiment was repeated using at least two biological replicates, and consistent results were confirmed by nsTEM.

## Acknowledgements

This work was supported by JSPS KAKENHI (grant nos. 19H02832, 19K22253, and 21H05116 to Yuta Suzuki (Y. Suzuki); 21H05117 to Y. Suzuki and Yukihiko Sugita (Y. Sugita); and 20K22628, 21J00530, and 22KJ1644 to M.N.), JST PRESTO (grant no. JPMJPR22A7 to Y. Suzuki.), Takeda Science Foundation to Y. Suzuki, Chubei Itoh Foundation to Y. Suzuki, and The Hakubi Center for Advanced Research to Y. Sugita and Y. Suzuki. Y. Suzuki also acknowledges additional support from the JST-CREST program (JPMJCR21B2; PI: Ryuji Kawano) for research activities related to the JST PRESTO project led by Y. Suzuki.

## Author contributions Statement

Y. Suzuki directed the project and led the conceptual framework. M.N. and Y. Suzuki conceived and designed the study. M.N. performed all experiments except cryo-EM and analyzed the data with Y.Suzuki. providing interpretative guidance. T.F., Y. Sugita and M.N. performed cryo-EM data collection, and T.F. and Y. Sugita analyzed the cryo-EM data. Y.Suzuki and M.N. wrote the manuscript with contributions from T.F. and Y. Sugita.

