## Supplementary Figures and Tables for "Thermally Tunable Heterodimeric Linkers Control Protein Tube Morphology"

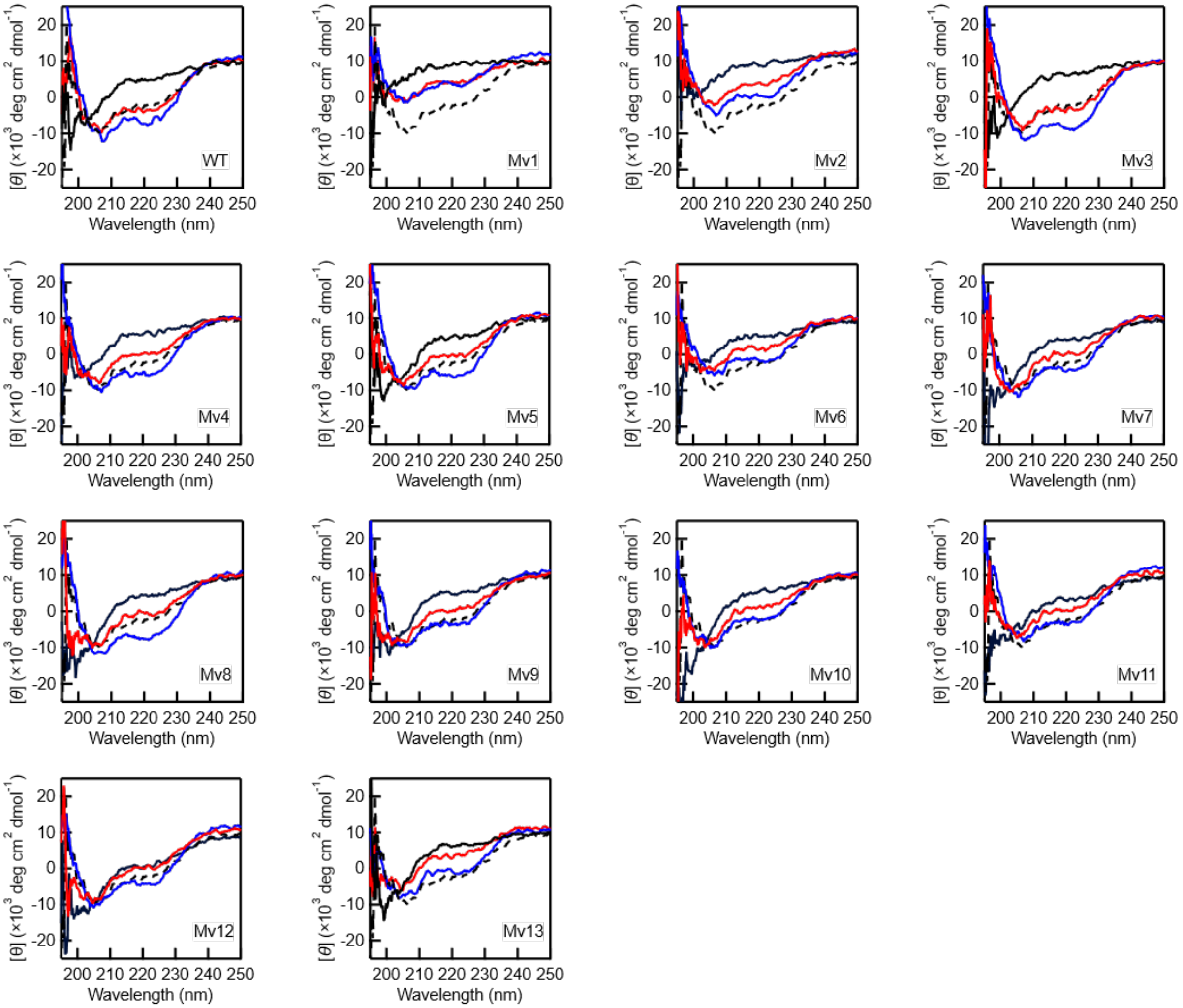

**Supplementary Fig. 1. Coiled-coil formation of M3L2(WT) and Mvs. |** CD spectra for isolated M3L2 variants (solid black lines) and p66α (dashed black lines) measured at 25 °C, as well as spectra for mixtures of each variant with p66α measured at 5 °C and 25 °C (blue and red lines, respectively). All spectra are labelled with the corresponding construct names. Source data are provided as a Source Data file.

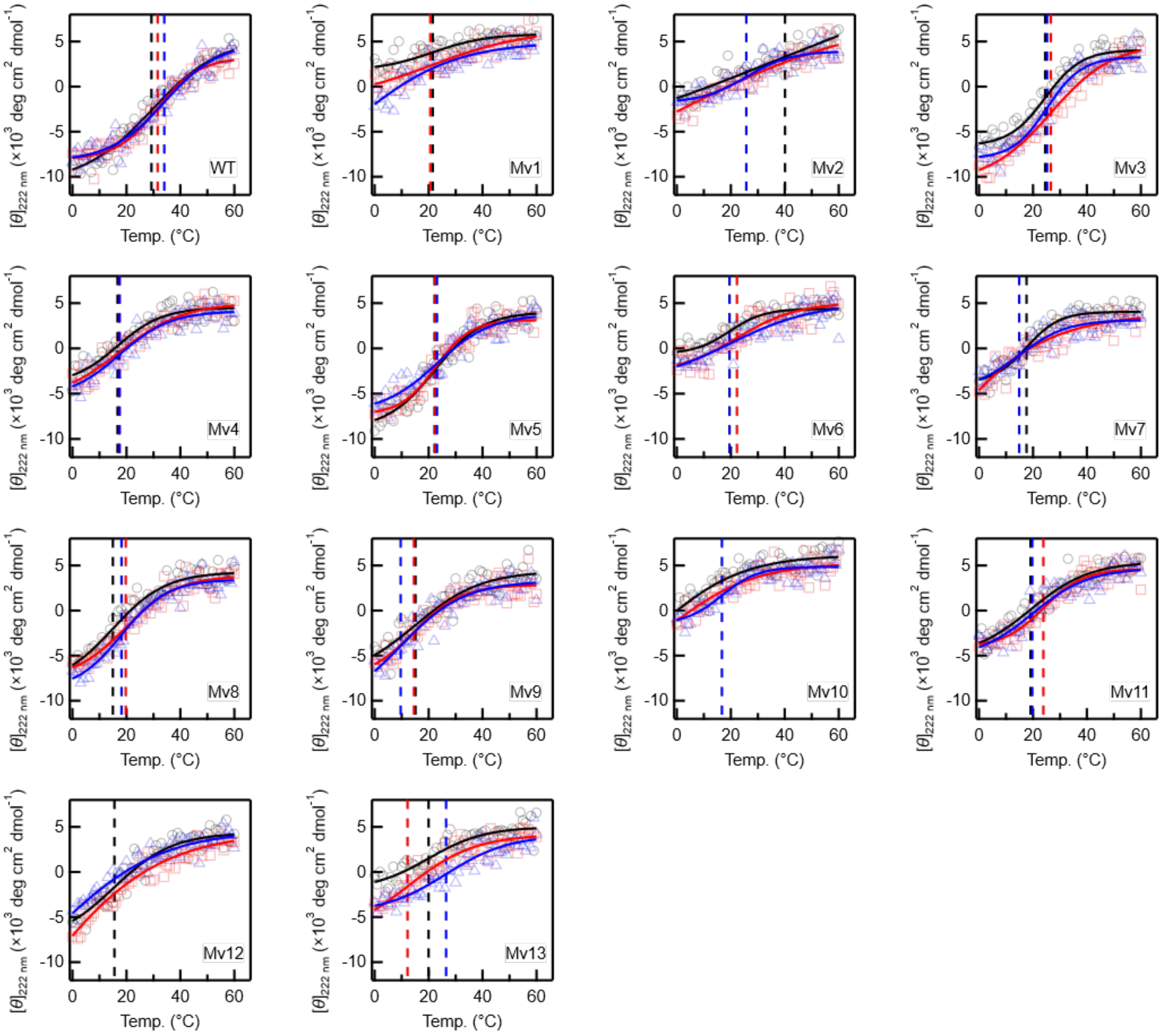

**Supplementary Fig. 2. Thermal stability of M3L2(WT) and Mvs.** | Thermal denaturation curves of mixtures of M3L2 variants with p66α recorded from 0 to 60 °C. For each variant, three independent experiments were performed; individual $\theta_{222\ \mathrm{nm}}$ values are plotted with different markers. Fitted curves (solid lines) and the extracted $T_m$ values (dashed lines) are shown in matching colours for each dataset. Numerical $T_m$ and the calculated $T_{m,Ave}$ values are summarised in **Supplementary Table 1**. Source data are provided as a Source Data file.

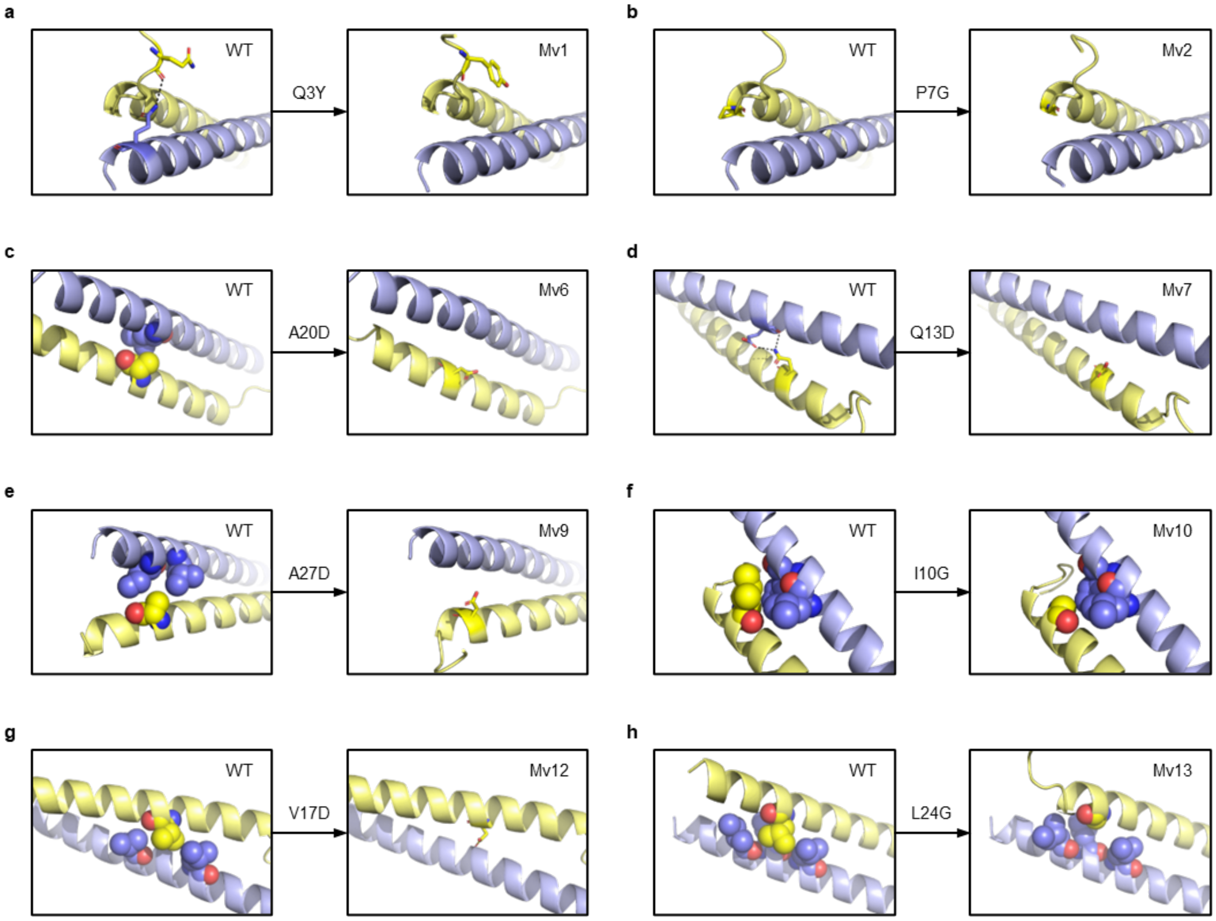

**Supplementary Fig. 3. AF3 structural models of Mvs/p66α coiled-coil complexes. | a–h,** the most reliable AF3-predicted models of WT (left) and each Mv variant (right) in complex with p66α, excluding the models shown in Fig. 2. For each model, the region surrounding the mutated residue is enlarged. Mutated residues and interacting partners are shown in ball (space-filling) or stick representations, with interchain hydrogen bonds indicated by dashed lines.

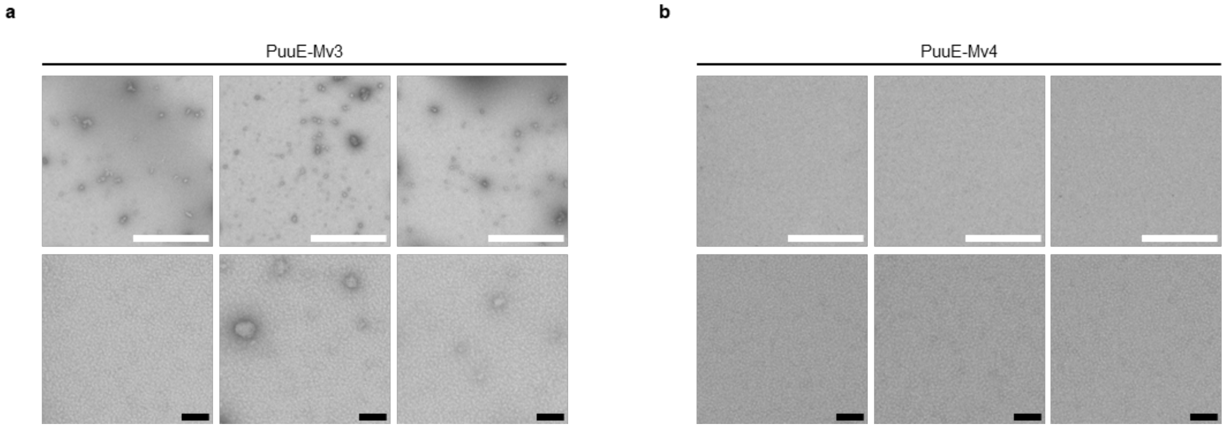

**Supplementary Fig. 4. nsTEM images of PuuE-Mvs in the absence of PuuE-p. | a, b,** Representative nsTEM images of PuuE-Mv3 (**a**) and PuuE-Mv4 (**b**). 12.5 μM PuuE-Mvs in experimental buffer was incubated at 25 °C for 24 h and imaged via nsTEM. Scale bars, 1 μm (white), 100 nm (black).

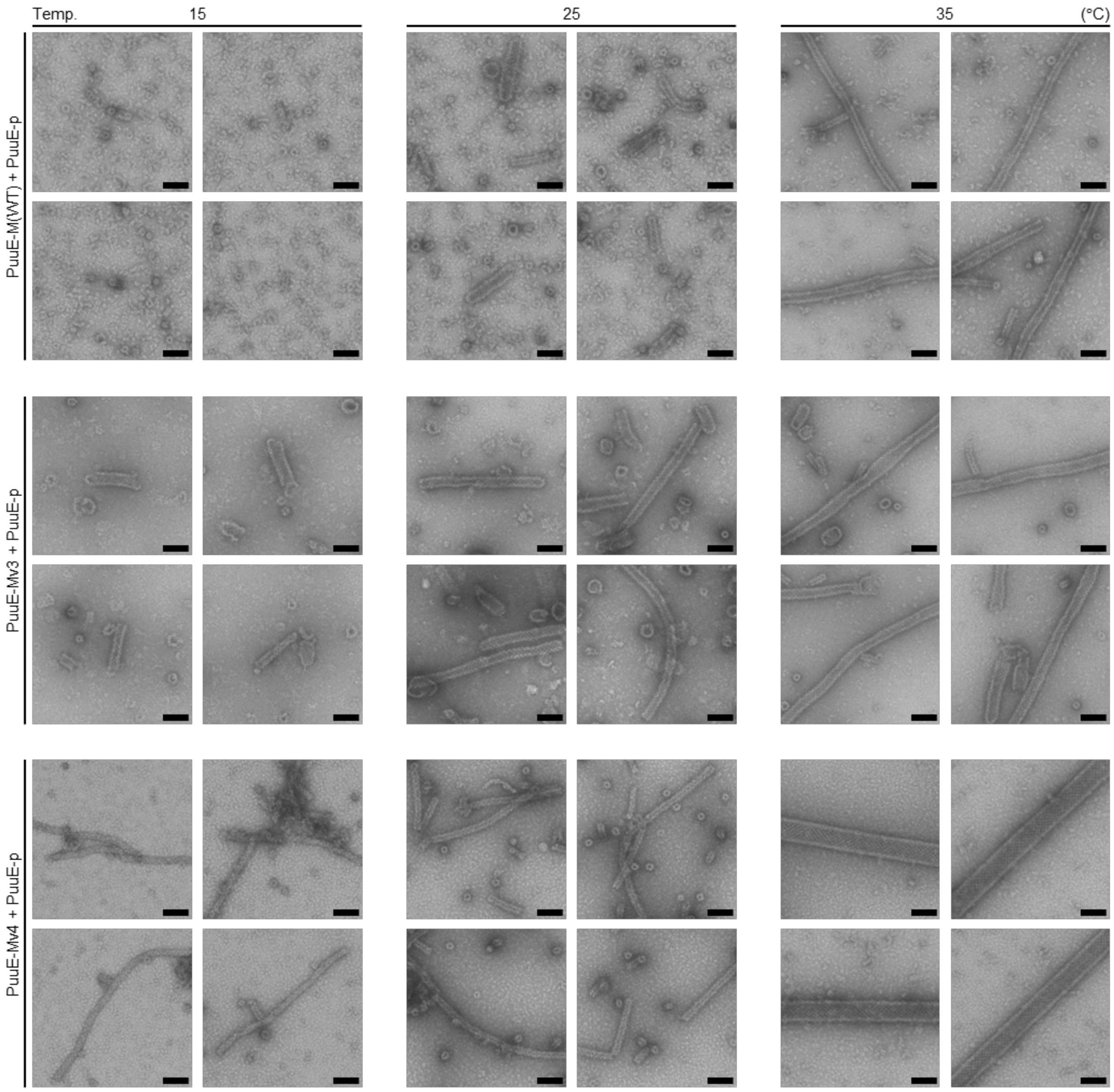

**Supplementary Fig. 5. Additional nsTEM images corresponding to Fig. 3b. |** Representative nsTEM images of tubes formed from PuuE-M(WT) (top), PuuE-Mv3 (middle), and PuuE-Mv4 (bottom) mixed individually with PuuE-p and incubated for 24 h under the temperature conditions indicated at the top (15, 25, and 35 °C).

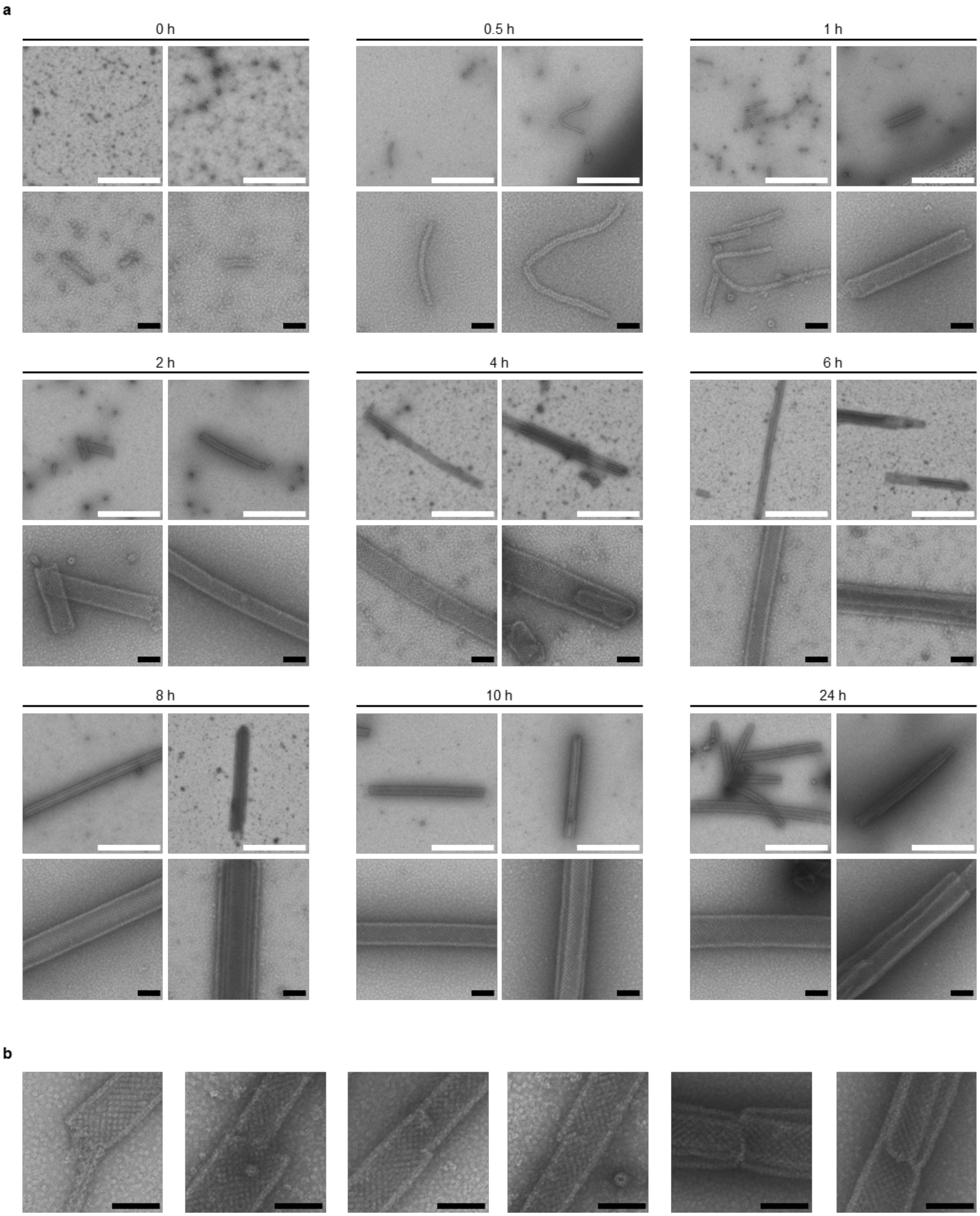

**Supplementary Fig. 6. nsTEM analysis of PuuE-Mv4 tube formation at 35 °C. | a,** Representative nsTEM images of tubes formed from PuuE-Mv4 mixed with PuuE-p and incubated at 35 °C from 0 to 24 h. These images provide additional time-course data complementary to Fig. 4 but show fields of view not included in the main figure. Time points are indicated at the top of each image. **b,** Representative nsTEM images showing junctions between tubes of different diameters and displaced tube-wall segments, captured in the same dataset as panel **a**. Scale bars: 1 µm (white); 100 nm (black).

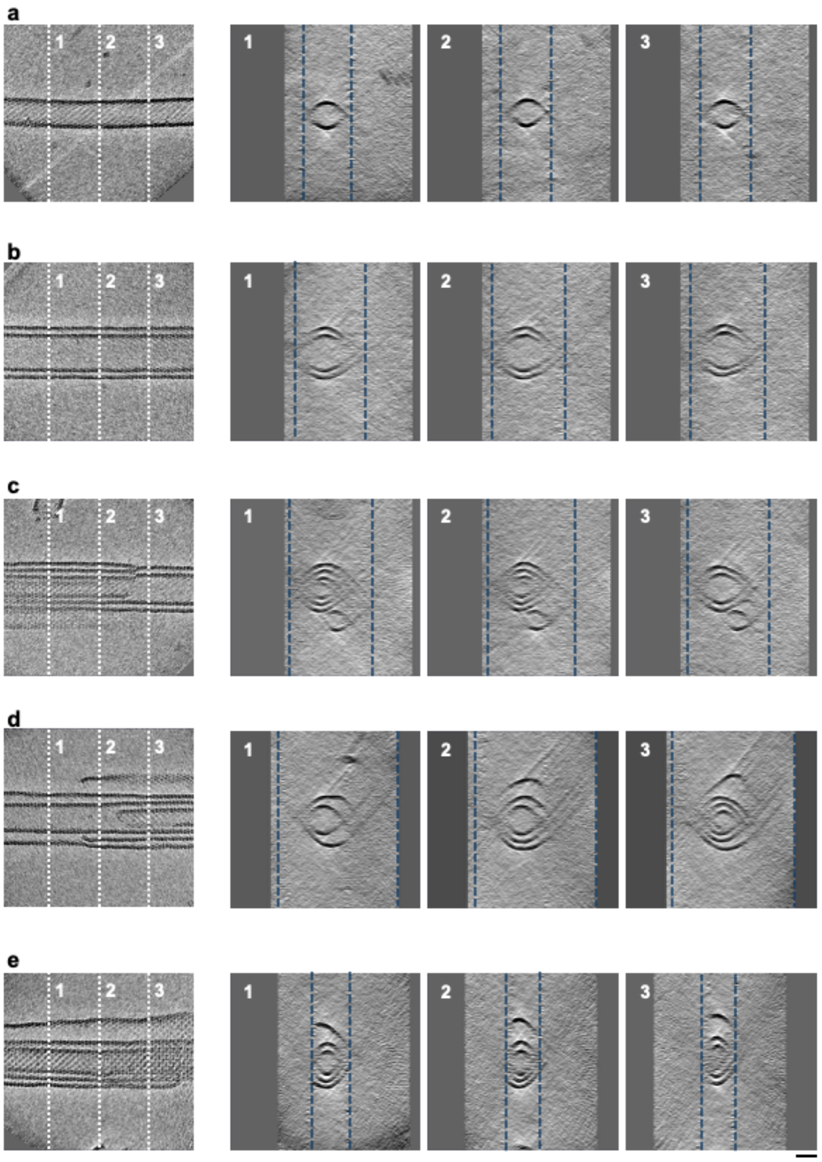

**Supplementary Fig. 7. Cryo-ET analysis of PuuE-Mv4 tube formation at 35 °C. | a–d**, Representative tomographic slices showing longitudinal views (left panel; single slice) and cross-sectional views (right panels; sum of 25 consecutive slices) of PuuE-Mv4 tubes in thicker ice (~120–280 nm) from cryo-ET reconstructions. Single- (a), double- (b), triple- (c), and four-layered (d) tubes are shown. **e**, PuuE-Mv4 tubes in ~80 nm of thin ice, showing pronounced tube flattening and an incomplete outer layer. White and blue dashed lines indicate the positions of cross-sections, labelled with numbers, and the ice surface, respectively. Scale bar: 50 nm.

### Supplementary Tables

| Name | Mutation | $\Delta\Delta G_{Pred}$ (kcal/mol) | $T_m$ (°C) | $T_{m,Ave}$ (°C) |
|---|---|---|---|---|
| WT | None | 0 | 29.3 ± 1.7 | 31.7 |
| | | | 31.6 ± 1.0 | |
| | | | 34.1 ± 1.6 | |
| Mv1 | Q3Y | 0.145 | 21.6 ± 5.8 | - |
| | | | 20.7 ± 13.3 | |
| | | | −5.8 ± 48.9 | |
| Mv2 | P7G | 0.489 | 40.1 ± 106.0 | - |
| | | | −4.5 ± 134.0 | |
| | | | 25.8 ± 2.4 | |
| Mv3 | A14G | 0.888 | 24.5 ± 1.1 | 25.5 |
| | | | 26.7 ± 1.4 | |
| | | | 25.3 ± 0.9 | |
| Mv4 | R16G | 1.136 | 16.7 ± 2.8 | 17.2 |
| | | | 17.5 ± 4.2 | |
| | | | 17.5 ± 3.1 | |
| Mv5 | L28G | 1.443 | 22.2 ± 1.6 | 22.6 |
| | | | 22.5 ± 1.0 | |
| | | | 23.1 ± 2.2 | |
| Mv6 | A20D | 1.476 | 19.5 ± 1.8 | - |
| | | | 22.3 ± 4.7 | |
| | | | 19.5 ± 9.9 | |
| Mv7 | Q13D | 1.546 | 17.6 ± 1.4 | - |
| | | | −21.6 ± 93.1 | |
| | | | 14.9 ± 4.1 | |
| Mv8 | R21D | 1.637 | 15.0 ± 3.2 | 17.7 |
| | | | 19.8 ± 2.2 | |
| | | | 18.2 ± 1.6 | |
| Mv9 | A27D | 1.759 | 15.2 ± 6.4 | - |
| | | | 14.6 ± 3.3 | |
| | | | 9.7 ± 7.5 | |
| Mv10 | I10G | 2.393 | −5.6 ± 44.0 | - |
| | | | −1.7 ± 34.8 | |
| | | | 16.7 ± 2.4 | |
| Mv11 | V5N | 2.439 | 19.1 ± 3.2 | 20.9 |
| | | | 23.9 ± 1.4 | |
| | | | 19.8 ± 3.4 | |
| Mv12 | V17D | 2.452 | 15.6 ± 4.2 | - |
| | | | −11.3 ± 45.1 | |
| | | | −4.6 ± 34.5 | |
| Mv13 | L24G | 2.802 | 20.0 ± 4.4 | - |

| | | | |
|---|---|---|---|
| | | | 12.2 ± 7.5 |
| | | | 26.5 ± 3.0 |

**Supplementary Table 1. M3L2(WT) and its mutants. |** Predicted stability changes for point mutations ($\Delta\Delta G_{Pred}$) obtained using ThermoMPNN are shown for M3L2(WT) and its mutants (Mvs). $T_m$ and $T_{m,Ave}$ determined from CD measurements are provided for coiled-coil complexes formed between p66α and each variant. $T_m$ values are shown with standard deviation. Source data are provided as a Source Data file.

**M3L2-Gly$_{50}$-p66α;** M3L2-Gly$_{50}$-p66α
SGQLVTPADIRRQARRVKKARERLAKALQADRLAGGGGGGGGGGGGGGGGGGGGGGGGGGG
GGGGGGGGGGGGGGGGGGGGGGGPEERERMIKQLKEELRLEEAKLVLLKKLRQSQIQ

**6xHis-TEVcs-M3L2(WT);** 6xHis-TEVcs-M3L2
MHHHHHHENLYFQGSGQLVTPADIRRQARRVKKARERLAKALQADRLA

**p66α;** 6xHis-TEVcs-p66α
MHHHHHHENLYFQGPEERERMIKQLKEELRLEEAKLVLLKKLRQSQIQ

**8xHis-TEVcs-M3L2(WT);** 8xHis-TEVcs-M3L2
MHHHHHHHHENLYFQGSGQLVTPADIRRQARRVKKARERLAKALQADRLA

**Mv1;** 8xHis-TEVcs-M3L2(Q3Y)
MHHHHHHHHENLYFQGSGYLVTPADIRRQARRVKKARERLAKALQADRLA

**Mv2;** 8xHis-TEVcs-M3L2(P7G)
MHHHHHHHHENLYFQGSGQLVTGADIRRQARRVKKARERLAKALQADRLA

**Mv3;** 8xHis-TEVcs-M3L2(A14G)
MHHHHHHHHENLYFQGSGQLVTPADIRRQGRRVKKARERLAKALQADRLA

**Mv4;** 8xHis-TEVcs-M3L2(R16G)
MHHHHHHHHENLYFQGSGQLVTPADIRRQARGVKKARERLAKALQADRLA

**Mv5;** 8xHis-TEVcs-M3L2(L28G)
MHHHHHHHHENLYFQGSGQLVTPADIRRQARRVKKARERLAKAGQADRLA

**Mv6;** 8xHis-TEVcs-M3L2(A20D)
MHHHHHHHHENLYFQGSGQLVTPADIRRQARRVKKDRERLAKALQADRLA

**Mv7;** 8xHis-TEVcs-M3L2(Q13D)
MHHHHHHHHENLYFQGSGQLVTPADIRRDARRVKKARERLAKALQADRLA

**Mv8;** 8xHis-TEVcs-M3L2(R21D)
MHHHHHHHHENLYFQGSGQLVTPADIRRQARRVKKADERLAKALQADRLA

**Mv9;** 8xHis-TEVcs-M3L2(A27D)
MHHHHHHHHENLYFQGSGQLVTPADIRRQARRVKKARERLAKDLQADRLA

**Mv10;** 8xHis-TEVcs-M3L2(I10G)
MHHHHHHHHENLYFQGSGQLVTPADGRRQARRVKKARERLAKALQADRLA

**Mv11;** 8xHis-TEVcs-M3L2(V5N)
MHHHHHHHHENLYFQGSGQLNTPADIRRQARRVKKARERLAKALQADRLA

**Mv12;** 8xHis-TEVcs-M3L2(V17D)
MHHHHHHHHENLYFQGSGQLVTPADIRRQARRDKKARERLAKALQADRLA

**Mv13;** 8xHis-TEVcs-M3L2(L24G)
MHHHHHHHHENLYFQGSGQLVTPADIRRQARRVKKARERGAKALQADRLA

**PuuE-M(WT);** 6xHis-TEVcs-PuuE-M3L2
MHHHHHHENLYFQGVDYPRDLIGYGSNPPHPHWPGKARIALSFVLNYEEGGERNILHGDKESEA
FLSEMVSAQPLQGERNMSMESLYEYGSRAGVWRILKLFKAFDIPLTIFAVAMAAQRHPDVIRAMV
AAGHEICSHGYRWIDYQYMDEAQEREHMLEAIRILTELTGERPLGWYTGRTGPNTRRLVMEEGG
FLYDCDTYDDDLPYWEPNNPTGKPHLVIPYTLDTNDMRFTQVQGFNKGDDFFEYLKDAFDVLYA
EGAEAPKMLSIGLHCRLIGRPARLAALQRFIEYAKSHEQVWFTRRVDIARHWHATHPYTSGQLVT
PADIRRQARRVKKARERLAKALQADRLA

**PuuE-p;** 6xHis-TEVcs-PuuE-p66α
MHHHHHHENLYFQGVDYPRDLIGYGSNPPHPHWPGKARIALSFVLNYEEGGERNILHGDKESEA
FLSEMVSAQPLQGERNMSMESLYEYGSRAGVWRILKLFKAFDIPLTIFAVAMAAQRHPDVIRAMV
AAGHEICSHGYRWIDYQYMDEAQEREHMLEAIRILTELTGERPLGWYTGRTGPNTRRLVMEEGG
FLYDCDTYDDDLPYWEPNNPTGKPHLVIPYTLDTNDMRFTQVQGFNKGDDFFEYLKDAFDVLYA
EGAEAPKMLSIGLHCRLIGRPARLAALQRFIEYAKSHEQVWFTRRVDIARHWHATHPYTPEERER
MIKQLKEELRLEEAKLVLLKKLRQSQIQ

**PuuE-Mv3;** 6xHis-TEVcs-PuuE-M3L2(A14G)
MHHHHHHENLYFQGVDYPRDLIGYGSNPPHPHWPGKARIALSFVLNYEEGGERNILHGDKESEA
FLSEMVSAQPLQGERNMSMESLYEYGSRAGVWRILKLFKAFDIPLTIFAVAMAAQRHPDVIRAMV
AAGHEICSHGYRWIDYQYMDEAQEREHMLEAIRILTELTGERPLGWYTGRTGPNTRRLVMEEGG
FLYDCDTYDDDLPYWEPNNPTGKPHLVIPYTLDTNDMRFTQVQGFNKGDDFFEYLKDAFDVLYA
EGAEAPKMLSIGLHCRLIGRPARLAALQRFIEYAKSHEQVWFTRRVDIARHWHATHPYTSGQLVT
PADIRRQGRRVKKARERLAKALQADRLA

**PuuE-Mv4;** 6xHis-TEVcs-PuuE-M3L2(R16G)
MHHHHHHENLYFQGVDYPRDLIGYGSNPPHPHWPGKARIALSFVLNYEEGGERNILHGDKESEA
FLSEMVSAQPLQGERNMSMESLYEYGSRAGVWRILKLFKAFDIPLTIFAVAMAAQRHPDVIRAMV
AAGHEICSHGYRWIDYQYMDEAQEREHMLEAIRILTELTGERPLGWYTGRTGPNTRRLVMEEGG
FLYDCDTYDDDLPYWEPNNPTGKPHLVIPYTLDTNDMRFTQVQGFNKGDDFFEYLKDAFDVLYA
EGAEAPKMLSIGLHCRLIGRPARLAALQRFIEYAKSHEQVWFTRRVDIARHWHATHPYTSGQLVT
PADIRRQARGVKKARERLAKALQADRLA

**Supplementary Table 2. Amino acid sequences of the protein constructs used in this study.**
